# Synaptic transmission induces site-specific changes in sialylation on *N*-linked glycoproteins in rat nerve terminals

**DOI:** 10.1101/2020.04.06.027524

**Authors:** Inga Boll, Pia Jensen, Veit Schwämmle, Martin R. Larsen

**Author notes:** Corresponding author: Prof. Dr. Martin R. Larsen: Department of Biochemistry and Molecular Biology, the University of Southern Denmark, Denmark.

## Abstract

Synaptic transmission leading to release of neurotransmitters in the nervous system is a fast and highly dynamic process. Previously, protein interaction and phosphorylation have been thought to be the main regulators of synaptic transmission. Here we show a novel potential modulator of synaptic transmission, sialylation of *N*-linked glycosylation. The negatively charged sialic acids can be modulated, similarly to phosphorylation, by the action of sialyltransferases and sialidases thereby changing local structure and function of membrane glycoproteins. We characterized site-specific alteration in sialylation on *N-*linked glycoproteins in isolated rat nerve terminals after brief depolarization using quantitative sialiomics. We identified 1965 formerly sialylated *N-*linked glycosites in synaptic proteins and found that the abundances of 430 glycosites changed after five seconds depolarization. We observed changes on essential synaptic proteins such as synaptic vesicle proteins, ion channels and transporters, neurotransmitter receptors and cell adhesion molecules. This study is to our knowledge the first to describe ultra-fast site-specific modulation of the sialiome after brief stimulation of a biological system.

## Introduction

Neurotransmission facilitates brain function by enabling a fast cell-cell signalling between neurons. In chemical synapses the electric potential is converted to a chemical signal by the influx of calcium and the subsequent exocytosis of neurotransmitters contained in synaptic vesicles in nerve terminals. The whole machinery of synaptic transmission includes several processes such as ion channel opening and closing, exocytosis and endocytosis as well as a diversity of receptor-ligand interactions and signalling cascades. These processes are of particular importance for diverse pathological processes in the brain, since several neurological diseases such as Alzheimer’s disease, Parkinson’s disease, autism, schizophrenia and epilepsy have been shown to have their main impairments in nerve terminals^1,2^. The molecular processes of synaptic transmission have been investigated extensively by using preparations of nerve terminals, the so-called synaptosomes, which represent a versatile model system of synapses, because they are metabolically intact, thereby allowing various stimulations leading to synaptic transmission^3,4^.

Synaptosomal preparations and the development of fast and sensitive mass spectrometry^5^ enabled the characterization of the proteome and phosphoproteome of nerve terminals^6–12^. The combination of mass spectrometry and imaging techniques further provided a comprehensive three-dimensional protein map of nerve terminals^13,14^. In addition to the studies aiming at characterizing naïve nerve terminals, investigating changes in post-translational modifications (PTMs) upon stimulation of neural activity is of high interest. PTMs are the addition of chemical groups or small proteins to proteins by enzymes in order to change protein activity, function and interaction. Especially in nerve terminals, protein regulation must be fast and reversible to accomplish the fast nature of the synaptic transmission process. Quantitative phosphoproteomics of resting and KCl-activated synaptosomes demonstrated changes in phosphorylation related with the basic machinery of synaptic transmission^11,12,15^. Nevertheless, changes in one PTM are likely to affect other PTMs building a whole network of PTM-crosstalk^16–18^. One of the other PTMs besides phosphorylation that can modify proteins and change the conformation and charge of a protein locally is sialylation of *N-*linked glycoproteins on the cell surface or within the secretory pathway. The attachment of a glycan to a polypeptide chain represents one of the most complex modifications in nature, since the glycan can vary in the combination of monosaccharides and in branching and linkages^19^. Sialylation, referring to the attachment of sialic acids, modifies the terminal branches of glycans^20^. This modification is omnipresent in the brain on glycoproteins and glycolipids, whereof gangliosides, sialylated glycosphingolipids, are the most abundant sialoglycoconjugates^21^. However, sialylated *N-* linked glycosylation of proteins in the brain plays a role in neural development and neurotransmission (reviewed in ^22^) and especially polysialylation, referring to the linkage of up to hundreds of sialic acids at one non-reducing end of a glycan, has been shown to be essential for the brain development even though only a few proteins have so far been observed to carry this large modification^23^.

Targeting sialoglycoconjugates of the brain by the application of exogenous sialidase and sialidase inhibitors revealed that sialylation can control neuronal and network excitability^24^. Moreover, St3gal2/3 double-knockout mice lacking the sialyltransferase responsible for sialylation of gangliosides and some glycoproteins showed dysmyelination, reduction of neuronal markers and cognitive disability^25^.

Focusing on *N-*linked glycosylation, sialic acid linkage and expression levels are changing during mouse brain development^26,27^. Furthermore, desialylation of cerebral glycoproteins was observed in neuroinflammation contributing to pathophysiological conditions^28^. These findings of changes in sialylated glycosylation were further supported by *in vivo* labelling of sialoglycans in the brain indicating that sialoglycoconjugates are concentrated at the synapse and their sialylation is “dynamic”, within a time frame of six hours, whereas the turnover of sialylated glycoproteins varies between different brain regions^29^.

Apart from this rather slow variability of sialylated glycosylation which could still be accomplished in the endoplasmic reticulum (ER) or Golgi apparatus some studies suggested that glycoproteins can also be modified directly at the synaptic junction^30,31^. With regard to this, a desialylation/sialylation cycle of gangliosides at the synapse has been proposed^32^. An altered sialylation directly at the synaptic cell surface could suggest a direct role in the modulation of synaptic transmission.

Interestingly, the function of the voltage-gated potassium channel Kv1.1 depends on its sialylation. This was studied in by transfecting the rat brain Kv1.1 cDNA into a Chinese hamster ovary glycosylation deficient cell line (Lec mutant) or by treating wildtype cells with sialidase^33^. It was hypothesized that the negative charge of sialic acids alters the surface potential of the ion channel and affects its gating machinery^34^. Not only voltage-gated ion channels, but also ligand-gated ion channels such as the nicotinic acetylcholine receptor (AcChoR) contain important sialylated *N-*linked glycans which modulate the channel conductance by attracting cations due to their negative charge^35^. In addition to this, transporters such as the GABA transporter GAT1 showed a reduced activity upon sialic acid removal^36^. Surface sialic acids bind Na^+^ ions rendering the function of Na^+^-dependent transporters^36^. This was not only revealed for GAT1 but also for amino acid neurotransmitter transporters^37^. Except from the attraction of cations, sialylation might also modulate protein conformation and protein-protein interactions regulating the process of neurotransmission^22^.

However, all conclusions for the involvement of sialic acids and glycosylation in processes involved in neurotransmitter release presented above, are made based on genetic removal of glycosylation sites (glycosites), glycosylation deficient mutants or treatment with sialidasesn which in all cases do not provide direct site-specific evidence for the involvement of a highly dynamic modulation of sialylation during brief induction of synaptic transmission. In addition, the time period of the studies presented above are hours or days, allowing the generation of newly synthesized proteins and not modulation of glycosylation.

Despite several studies mapping *N-* and *O*-linked glycosylation in isolated synaptosomes^38,39^ and mouse brain tissue^27,40^, there is no information regarding the regulation of sialylation on surface proteins in conjunction with synaptic transmission in synaptosomes.

The present study aimed at identifying sialylated *N-*linked glycosites in rat synaptosomes and at the same time quantifying site-specific changes in sialylated glycosylation upon brief depolarization of synaptosomes. Here, we revealed for the first time the sialiome (defined as the large scale characterization of sialic acid containing glycosylation sites)^41^ of nerve terminals and identified site-specific changes in sialylation on selected proteins upon synaptic transmission. Despite the presumable lack of ER and Golgi in isolated synaptosomes, we managed to identify several sialyltransferases in the enriched active zone of nerve terminals that could explain the increased sialylation on some *N-*linked glycosites observed in this study. Our results clearly show that site-specific dynamic regulation of sialylation on synaptic proteins, including various ion-channels, ion-transporters, receptors and surface adhesion molecules, could be important for synaptic transmission. This is to our knowledge the first study on global site-specific changes in sialylation on *N*-linked glycoproteins upon brief stimulation of any biological system showing that sialylation can be as dynamic as well-known dynamic PTMs such as phosphorylation.

## Experimental Procedures

All chemicals were obtained from Sigma Aldrich, if not indicated otherwise.

### Synaptosome preparation and stimulation

Eight to ten weeks old male Sprague Dawley rats (Taconic) were decapitated unanesthetized and their brains were immediately dissected and homogenized in 0.32 M sucrose, 1 mM EDTA, 5 mM Tris base, pH 7.4 following a standard Percoll density gradient procedure^3^. After loading on a Percoll density gradient, only the F4 layer of the gradient was used for subsequent stimulation, since it contained the purest synaptosomes, however in minute amounts. Synaptosomes were resuspended in HEPES-buffered Krebs-like buffer (HBK) (118 mM NaCl, 4.7 mM KCl, 1.18 mM MgSO_4_, 0.1 mM Na_2_HPO_4_, 1.2 mM CaCl_2_, 25 mM NaHCO_3_, 10 mM glucose, 20 mM HEPES, pH 7.4) and incubated for 1 h at 37 °C to achieve a metabolic equilibrium. The equilibrated synaptosomes fraction was aliquoted to several aliquots each containing ca. 500 µg synaptosomal protein. For stimulation, the addition of an equal volume of high KCl HBK (147.7 mM KCl, 1.18 mM MgSO_4_, 0.1 mM Na_2_HPO_4_, 1.2 mM CaCl_2_, 10 mM glucose, 20 mM HEPES, pH 7.4) led to an increase in the KCl concentration to 76.2 mM inducing the depolarization of the membrane. Stimulation with HBK with 4.7 mM KCl served as a control. After 5 s, the stimulation was quenched by adding hot lysis buffer (final concentrations: 0.1% SDS (GE Healthcare), 25 mM HEPES, Complete protease inhibitor cocktail (Roche), PhosStop phosphatase inhibitor (Roche)) and immediate boiling at 95 °C for 8 min. The boiled samples were snap frozen in liquid nitrogen and stored at -80 °C until further usage.

### Ultracentrifugation and protein reduction, alkylation and digestion

Since the synaptosomal membrane was only partially dissolved in 0.1% SDS, an SDS-insoluble fraction was separated from the SDS-soluble proteins by ultracentrifugation for 90 min at 100,000 g. The SDS-insoluble fraction was solubilized by adding 6 M urea, 2 M thiourea, 10 mM dithiotreitol (DTT) and incubating for 30 min at RT. The reduced proteins were alkylated for 30 min in the dark by adding 20 mM iodoacetamide (IAA). For efficient trypsin digestion, the proteins were incubated for 2 h with 1 µl Lys-C (Wako). Subsequently, the solution was diluted ten times with 20 mM triethylammonium bicarbonate (TEAB), pH 7.6 prior to probe sonication for 2×10 s on ice. The protein concentration was determined using the Qubit™Protein Assay Kit (Thermo Fisher Scientific) and 100 µg protein of each sample were transferred to a new tube and digested overnight using 2 µg trypsin (Sigma, purified and methylated in-house). After digestion, the peptides were lyophilized. The SDS-soluble fraction was loaded on an Amicon Ultra Centrifugal Filter 10 K (Merck Millipore). After reduction of the volume by centrifugation at 10000 g for 40 min, 6 M urea, 2 M thiourea, 10 mM DTT were added and incubated for 30 min at RT. After incubation, the volume was reduced by centrifugation and protein alkylation was performed by adding 20 mM IAA and incubating for 30 min at RT. The proteins were washed twice with 20 mM TEAB, pH 7.6 prior to protein quantitation by Qubit as described above. For digestion overnight, 2% (w/w) trypsin were added to the proteins in the filter. After digestion, the solution in the filter was transferred to a low-binding tube and the filter was washed with 50 % acetonitrile (ACN). A total of 100 µg peptides from each replicate were transferred to another tube and lyophilized for subsequent analysis.

### Isobaric peptide labelling for relative quantitation

A total of 100 µg peptides from each sample were labeled using the TMT11plex Mass Tag Label Kit (Thermo Scientific) according to the manufacturer’s instructions. Briefly, peptides were dissolved in 75 µl 100 mM TEAB, pH 8.3 and mixed with 0.5 mg TMT reagent dissolved in 25 µl anhydrous ACN. After incubation for 1 h, efficient labelling and equal protein amounts were checked by LC-MS/MS analysis. The labelling reaction was quenched by adding 0.54 µl 50 % hydroxylamine (Merck) and incubating for 15 min. Subsequently, the samples were combined according to the LC-MS/MS test run and lyophilized.

### Enrichment of formerly sialylated N-linked glycopeptides

The enrichment of sialylated *N-*linked glycopeptides was performed by using Titanium Dioxide (TiO_2_) chromatography that we previously have shown to be more than 95% selective for sialylated *N-*linked glycopeptides41. Here we combined the TiO_2_ in a modified version of the TiSH protocol, we previously published42. The protocol uses an initial TiO_2_ enrichment step to separate phosphopeptides and sialylated glycopeptides from all remaining peptides (hereafter called non-modified). After enzymatic deglycosylation, sequential elution from IMAC (SIMAC) and a second TiO2 enrichment further separated the formerly sialylated *N-* linked glycopeptides from the phosphorylated peptides^43^. Shortly, peptides were dissolved in loading buffer (1 M glycolic acid, 80 % ACN, 5 % trifluoroacetic acid (TFA)) incubated for 15 min with 0.6 mg TiO_2_ beads per 100 µg peptide solution (Titansphere, GL Sciences Inc) using vigorous shaking. After pelleting the beads, the supernatant was subjected to a second TiO_2_ incubation using half of the TiO_2_ beads. Subsequently to washing, phosphopeptides and sialylated glycopeptides were eluted for 15 min on a shaker with 1 % ammonia solution, pH 11.3. This elution was lyophilized, redissolved in 20 mM TEAB, pH 7.6 and incubated overnight with 2 µl PNGase F (New England BioLabs) and 0.5 µl sialidase A (Prozyme). After employing SIMAC for the enrichment of multi-phosphorylated peptides^43^, the flow-through and washes were subjected to a second TiO_2_ enrichment using 70 % ACN, 2 % TFA as a loading buffer. The peptides that did not bind to the TiO_2_ beads contained formerly sialylated *N-*linked glycopeptides and were lyophilized for subsequent analysis. The phosphopeptide elution was lyophilized as well and analyzed for evaluation of the stimulation effects.

### Micropurification

Peptides were micropurified by using an in-house protocol with custom-made microcolumns in p200 pipette tips. These were stowed with an Empore™ C18 Solid Phase Extraction Disk (Supelco) plug and packed with Oligo R3 Reversed Phase Resin (Applied Biosystems). Samples were acidified with 10 % TFA, 100 % formic acid or redissolved in 0.1% TFA. After equilibration of the column with 0.1 % TFA, samples were loaded slowly, and the column was washed with 0.1 % TFA. For elution, 60% ACN, 0.1% TFA were applied. Peptides were lyophilized prior to offline fractionation.

### High pH reversed-phase fractionation

To decrease the complexity, samples were dissolved in 20 mM ammonium formate, pH 9.3 (buffer A) and loaded onto an Acquity UPLC® M-Class CSH™ C18 column (Waters) for offline reversed-phase high pH fractionation using a Dionex Ultimate 3000 HPLC system (Thermo Scientific). Peptides were separated by increasing concentrations of buffer B (80 % ACN, 20 % buffer A) from 2 % to 50 % buffer B in 102 min and from 50 % to 95 % buffer B in 5 min with a flow-rate of 5 µl/min. The eluting peptides were collected in 20 concatenated fractions using 2 min elution per fraction and dried down prior to LC-MS/MS analysis. Peptides that precipitated in buffer A were run directly by LC-MS/MS without offline fractionation.

### Reversed phase nanoLC-ESI-MS/MS

All samples were dissolved in 0.1 % formic acid (FA) and analyzed by nanoLC-ESI-MS/MS using an EASY-nLC 1000 (Thermo Scientific) and a Q-Exactive HF mass spectrometer (Thermo Scientific). The in-house made fused silica capillary two-column setup consisted of a 3 cm pre-column with 100 µm inner diameter packed with Reprosil-Pur 120 C18-AQ, 5 µm (Dr. Maisch GmbH) and an 18 cm pulled emitter analytical column with 75 µm inner diameter packed with Reprosil-Pur 120 C18-AQ, 3 µm (Dr. Maisch GmbH). Peptides were separated by a gradient starting from 99 % buffer A (0.1 % FA), 1 % buffer B (95 % ACN, 0.1 % FA) and increasing to 3 % buffer B in 3 min. The gradient continued with a step from 3 % to 28 % buffer B in 50 min and from 28 % to 45 % buffer B in 8 min until elevating the concentration of buffer B to 100 % in 3 min. The nLC was used with a flowrate of 250 nl/min connected online to the mass spectrometer working in a data-dependent acquisition mode. The acquisition in positive ion mode comprised full MS scans from 400-1600 m/z with a resolution of 120,000 full width half maximum (FWHM) using a maximum filling time of 100 ms and an automatic gain control (AGC) target value of 3×10^6^ ions. Depending on the sample complexity, the 10 or 20 most intense ions were selected for HCD fragmentation with a normalized collision energy of 34. The MS2 spectrum was acquired from 110 to 2000 m/z with a resolution of 60,000 FWHM using a maximum filling time of 100 ms and an AGC target value of 1×10^5^ ions combined with an isolation window of 1.2 m/z as well as dynamic exclusion of 30 s.

### MS data searching

All raw files were searched against the Swissprot *mus musculus* and *rattus norvegicus* references (downloaded 18.12.2017) in Proteome Discoverer (PD) 2.3.0.520 (Thermo Scientific) using an in-house Mascot server combined with the Percolator for peptide validation (1 % FDR for proteins and peptides). We used the mouse database (25131 entries) to compensate proteins that were not covered by the rat database (9617 entries), selecting *rattus norvegicus* as a preferred taxonomy in PD. The search included the following parameters: static modifications: TMT6plex on lysine and the N-terminus as well as carbamidomethyl on cysteines; dynamic modifications: deamidation of asparagine or phosphorylation of serine, threonine and tyrosine; tryptic peptides with maximal one missed cleavage allowed; precursor mass tolerance 10 ppm, fragment mass tolerance 0.05 Da; Mascot ion score ≥ 20, only rank one peptides.

All non-modified datasets were first searched against the entire *mus musculus* and *rattus norvegicus* proteomes without considering any dynamic modifications. Afterwards, the spectra were further subjected to a second search using a custom-made database that contained all identified sialylated glycoproteins. Therefore, the identified sialylated glycoproteins were used for the creation of a fasta-file using the Swissprot database. In PD, the non-modified datasets were searched with Sequest HT as a search engine against these reduced glycoprotein databases filtering PSMs for an XCorr threshold ≥ 0.9. The results of this search were merged with the first search and used for abundance adjustment of the sialylated glycopeptides to their respective non-modified counterpart. All results from PD were exported as an Excel file and further processed in Excel (Microsoft Office).

### Further data processing

After performing the search in PD and exporting the normalized abundances, non-modified proteins were filtered keeping only the Master proteins that were identified with at least two unique peptides. These proteins were subjected to statistical testing or abundance adjustment of formerly sialylated *N*-linked glycopeptides.

Formerly sialylated *N-*linked glycopeptides were determined by searching for deamidation of asparagine and subsequent filtering for the Asn-Xaa-Ser/Thr/Cys *N-*glycosylation motif. Only peptides with one deamidation were considered and only membrane bound or secreted proteins were used for further analysis. Due to protein translocation between the SDS-insoluble and the SDS-soluble compartments, all formerly sialylated *N-*linked glycopeptides were adjusted to the expression of their respective non-modified protein abundance. Formerly sialylated *N-*linked glycopeptides without a respective non-modified protein identified could not be adjusted and are marked specifically in the following.

### Experimental design and statistical rationale

For each experiment two rat brains were pooled for the preparation of synaptosomes, whereof the isolated synaptosomes were aliquoted in order to subject at least four samples to the control or high KCl stimulation (n=4). Both fractions, SDS-soluble and SDS-insoluble were analyzed in separate TMT sets. The initial experiment (experiment 1) served as a quality control of the enrichment of formerly sialylated *N-*linked glycopeptides and only qualitative data were considered, whereas experiment 2 was analysed quantitatively as well. Formerly sialylated *N-* linked glycopeptides followed normal distribution and were considered as significantly changing, if q<0.05 after performing LIMMA testing and subsequent correction for multiple testing by Storey^44,45^. Furthermore, only log-ratios greater than 0.26 or smaller than -0.26 were considered as significantly changing.

### Bioinformatic analysis of significantly changing proteins

If necessary, peptide abundances were rolled-up with regard to proteins using Perseus^46^. Protein networks were analyzed using Cytoscape combined with the STRING application (confidence score 0.7) which also included enrichment analysis of KEGG pathways^47–49^. For enrichment analysis of proteins, DAVID was used defining all identified proteins or the proteome of *rattus norvegicus* as a background^50^. For enrichment analysis, q-values <0.05 after Benjamini-Hochberg correction were considered as significant. For the identification of PTM enzymes, all identified proteins were compared to all proteases, phosphatases, kinases and glycosyltransferases by using the QuickGO tool, searching for peptidase activity, phosphoprotein phosphatase activity, protein kinase activity and protein *N-*linked glycosylation, respectively. All histograms were made using Prism (GraphPad). Protein structures were modified and displayed in Pymol (Schrödinger). Schematic diagrams were produced with the help of Science Slides (VisiScience).

### Determination of glutamate release from synaptosomes

To investigate their intactness and metabolic activity, a small portion of the isolated synaptosomes was used to determine glutamate release. Synaptosomes were incubated for 10 min at 37 °C either with high KCl HBK or HBK followed by centrifugation for 15 s at 13400 g to pellet the synaptosomes. The supernatants containing glutamate released from synaptic vesicles were diluted and inactivated by the addition of 0.2 M HCl. After neutralization by adding 1 M Tris base, the glutamate content was determined with the Glutamate Glo Assay Kit (Promega). The assay was performed in a 384 well plate according to the manufacturer’s instructions. Briefly, the assay used enzymatic reactions to measure a luminescence proportional to the glutamate concentration. The relative luminescence was measured using a Fluostar Omega plate reader (BMG labtech).

## Results and discussion

### Strategy for site-specific assessment of N-linked sialylation in rat synaptosomes

The strategy for investigating alteration in the sialiome of synaptosomes upon brief depolarization is illustrated in Figure 1. In order to investigate sialylated *N-*linked protein glycosylation in synaptosomes, these were isolated from rat brains using Percoll gradients^3^, and then subsequently stimulated for five seconds with a high KCl concentration (76 mM) or a control buffer. The reaction was terminated using boiling SDS to a final concentration of 0.1 % and further incubated at 95 °C for 8 min before the proteins were separated into two fractions, 0.1 % SDS-insoluble and 0.1 % SDS-soluble (in the following denoted as SDS-insoluble and SDS-soluble), using ultracentrifugation. Proteins were digested to peptides using trypsin and the resulting peptides were labelled with isobaric tandem mass tags (TMT) prior to the enrichment of sialylated *N-*linked glycopeptides by titanium dioxide (TiO_2_) chromatography, which provide a selective enrichment of sialylated *N-*linked glycopeptides, when using a special loading buffer^41,51^. The flow-through from the TiO_2_ chromatography contained the “non-modified” peptides that were used for quantitative proteomics analysis. After deglycosylation, the deglycopeptide fraction and non-modified peptide fraction were fractionated using high pH reversed phase (RP) separation, and peptide fractions were analysed by LC-MS/MS analysis (Figure 1A). Proteins and formerly sialylated *N-*linked glycopeptides were identified in two separate experiments.

**Figure 1:**
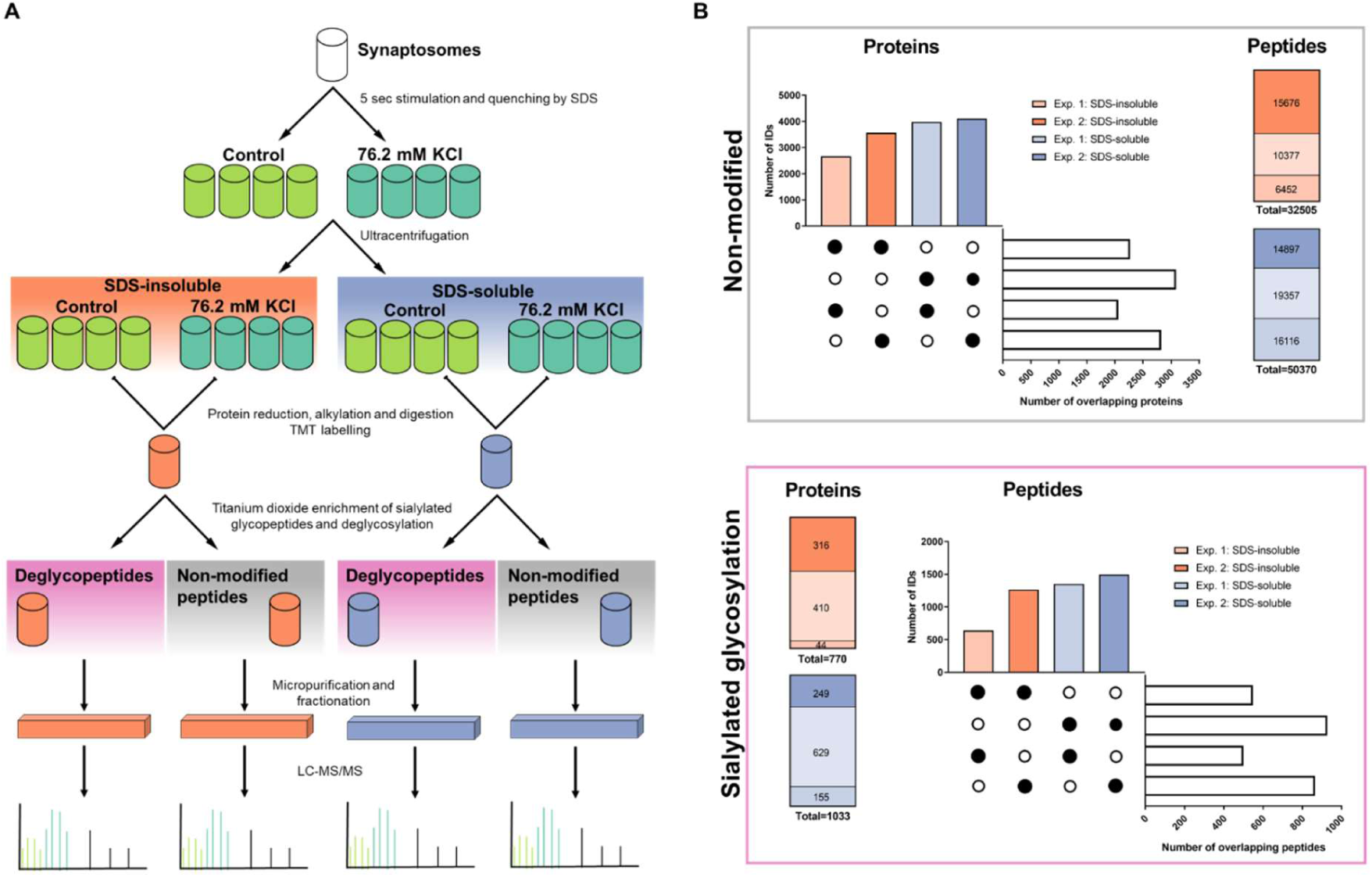
Summary of the experimental workflow and identified peptides and proteins. (A) Synaptosomes isolated from adult male wild-type Sprague-Dawley rats were depolarized for five seconds and subsequently quenched by the addition of boiling SDS lysis buffer. SDS-insoluble and SDS-soluble protein fractions were separated by ultracentrifugation followed by reduction, alkylation and digestion of proteins. Peptides were subjected to TMT labelling for relative quantification prior to the enrichment of sialylated glycopeptides by titanium dioxide. After enrichment and deglycosylation of sialylated glycopeptides, peptides were fractionated by high pH reversed-phase chromatography before analysing them by LC-MS/MS. (B) Identified proteins and peptides (FDR < 0.01) of the non-modified group and the enriched formerly sialylated *N-*linked group are shown. The identifications from two independent experiments as well as the overlap between the SDS-insoluble and SDS-soluble fractions and the two different experiments are compared.

### Overview of the synaptic proteome and sialiome

The strategy identified 3956 non-modified proteins (all peptides from the TiO_2_ flow-through) in the SDS-insoluble fraction in total in the two experiments (2270 proteins identified in both experiments) and 4998 non-modified proteins in the SDS-soluble fraction (3090 proteins identified in both experiments) (Figure 1B). Comparing only the identified proteins, both SDS fractions had an overlap of more than 80 % and in total 5692 proteins were identified. The SDS-insoluble fraction was highly enriched in active zone proteins such as protein Bassoon and Piccolo (Figure S1).

By examining sialylated *N-*linked glycoproteins, 770 and 1033 proteins were identified in the SDS-insoluble and SDS-soluble fraction, respectively. These proteins contained 2198 different formerly sialylated *N-*linked glycopeptides in both fractions and experiments together (1965 distinct *N-*linked glycosites). In general, the SDS-soluble fraction contained more formerly sialylated *N-*linked glycopeptides than the SDS-insoluble fraction. In the SDS-soluble fraction, a total of 1923 formerly sialylated *N-*linked glycopeptides were identified, whereas 1362 peptides were identified in the SDS-insoluble fraction. This is most likely due to the relative low amount of protein contained in the SDS-insoluble fraction, providing lower starting material prior to TiO_2_ chromatography and consequently a lower coverage of the sialiome. In general, the SDS-insoluble and SDS-soluble fraction were overlapping in about 50 % of the identified formerly sialylated *N-*linked glycopeptides.

For the first time, formerly sialylated *N-*linked glycosites in synaptosomes were determined exceeding the former knowledge about sialylated *N-*linked glycosylation in this compartment of neurons. Trinidad *et al*. investigated *N-* and O-glycosylation in murine synaptosomes and identified 298 intact sialylated *N-*linked glycopeptides, but these reflected only 10 individual peptides that were identified with different sialylated *N-*linked glycans^39^. The increased number of formerly sialylated *N-*linked glycosites identified in this study provides a starting point for further investigations on the individual protein level. The consideration of sialylated *N-*linked glycosites possessing the Asn-Xaa-Ser/Thr/Cys (Xaa#Pro) motif diminished the rate of false positives, however it needs to be noted that spontaneous deamidation can occur, especially in Asn-Gly-Ser/Thr/Cys and Asn-Ser-Ser/Thr/Cys consensus sites and further validation of these sites might be needed^52^. However, manual evaluation of a subset of identified sites using the Uniprot database containing known *N-*linked sites revealed a very low number of false positives.

### Synaptosome depolarization validation

To evaluate the depolarizing effect of the stimulation, phosphorylation sites of known synaptic transmission marker proteins and the release of the neurotransmitter glutamate were evaluated (Figure S2). Synaptosomes used for proteomics experiments released glutamate upon stimulation indicating an intact synaptosomal membrane and metabolic activity. Moreover, dephosphorylation of dynamin-1 on Ser-774 and Ser-778 as well as phosphorylation of synapsin-1 on Ser-566 were observed upon depolarization as described in the literature^53–55^. Since synaptic transmission is an ultra-fast process lasting only for milliseconds, depolarization for five seconds would include multiple synaptic transmission events and might induce artificial effects^56^. However, important features of synaptic transmission would still be measurable within this time frame.

### The sialiome roadmap of synaptosomes

By using isolated synaptosomes and further separating proteins into two fractions according to their SDS-solubility, obtaining an enriched active zone, a comprehensive map of sialylated *N-* linked glycosites was accomplished containing 1965 individual *N-*linked glycosites on 1139 proteins. Many proteins carrying sialylated *N-*linked glycosylation play an important role in synaptic transmission which was indicated by the significant enrichment of KEGG pathways such as neuroactive ligand-receptor interaction, axon guidance, glutamatergic synapse and calcium signalling pathways (Figure S3A). To reveal more specific functions, GO terms and KEGG pathways with a low corrected p-value and a high fold enrichment were examined further (Figure S3B). In the SDS-insoluble fraction, the identified sialylated *N-*linked glycoproteins belonged to ionotropic glutamate receptor signalling pathways, GABAergic and glutamatergic synaptic transmission and membrane depolarization processes. Similar functions were found for the glycoproteins identified in the SDS-soluble fraction. To reveal the differences between the SDS-insoluble and the SDS-soluble fraction regarding their unique sialylated glycoproteins, significantly enriched Uniprot keywords were compared (Figure 2A). Compared to the SDS-soluble fraction, the SDS-insoluble fraction showed a higher enrichment of ion channels and transporters, particularly for calcium, highly important for calcium signalling during synaptic transmission. Moreover, some synapse and postsynaptic sialylated membrane glycoproteins were exclusively identified in the SDS-insoluble fraction. This supports that the separation using SDS solubility is providing an enriched active zone fraction, as a small part of the postsynapse that is strongly bound to the active zone part of the presynapse is co-purified during the Percoll gradient separation of synaptosomes. In addition, these results show that the active zone in synaptosomes contains unique sialylated glycoproteins highly important for the function of the active zone, synaptic transmission. In contrast to this, the SDS-soluble fraction contained sialylated glycoproteins of lysosomal origin and proteins comprising EGF-like domains. Moreover, this fraction was enriched in calcium binding or calcium dependent proteins.

**Figure 2:**
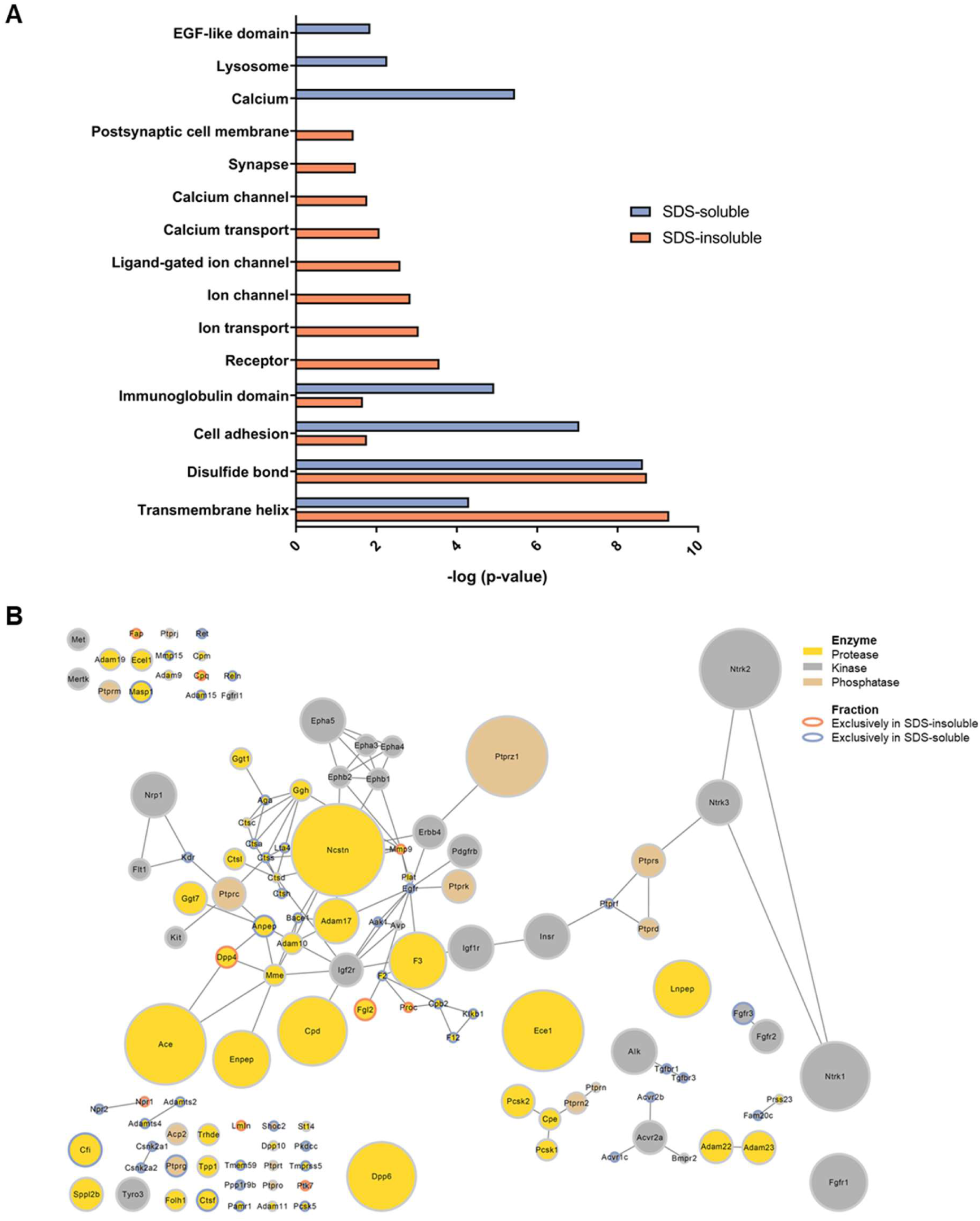
Characterization of the sialiome in the SDS-insoluble and SDS-soluble fraction. (A) Sialylated glycoproteins that were exclusively identified either in the SDS-insoluble or SDS-soluble fraction were analysed regarding their enrichment of Uniprot keywords. Key words with a Benjamini-Hochberg corrected p-value < 0.05 are displayed. (B) All identified formerly sialylated *N*-linked glycoproteins were screened for enzymes catalysing post-translational modifications. A network of identified proteases, kinases and phosphatases present in the SDS-insoluble, SDS-soluble or both fractions was created where the node size is proportional to the number of identified sialylated *N-*linked glycosites.

Apart from the differences between the two fractions, many proteins carrying sialylated *N-* linked glycosylation were kinases, phosphatases or proteases (Figure 2B). The group of proteases contained many members of the a disintegrin and metalloproteinase (ADAM) family, namely ADAM9, 10, 11, 17, 19, 22 and 23. In addition to this, the nicastrin subunit of the gamma-secretase was identified to carry sialylated *N-*linked glycans. This particular enrichment of proteases reflects the fact that glycosylation, and probably sialylation^57^, is important for protease activity^58^. For ADAMs, it is known that their enzymatic activity depends on their glycosylation and they perform activity-dependent proteolysis of synaptic cell adhesion molecules such as neuroligin-1^59–62^. Further studies investigating the impact of sialic acids on specific *N-*linked glycans in ADAM proteases in synaptosomes might be of high interest to shed light on their involvement in synaptic transmission and learning and memory related processes. Besides proteases, several kinases and phosphatases were identified to carry sialylated *N-*linked glycosylation, including kinases and phosphatases such as the receptor tyrosine-protein kinase erbB-4 (Erbb4) and the receptor-type tyrosine-protein phosphatase zeta (Ptprz1). These proteins were found to play a role in schizophrenia^63^. Other phosphatases of the receptor-type tyrosine-protein phosphatase family (Ptprc, Ptprn, Ptprn2, Ptprs) and prominent kinases (e.g. Ntrk1, Ntrk2, Ntrk3, Epha3, Epha5,) were present in the dataset of formerly sialylated *N-*linked glycoproteins indicating an impact of sialylated glycosylation on neuronal signalling. The relative high amount of important proteins for synaptic transmission in the synaptosomes that carried sialylated *N-*linked glycosylation, including enzymes for modulation of other PTMs, raised the idea to investigate, if sialylation can be dynamically modulated during synaptic transmission. Considering the lack of an ER and only vague evidence of Golgi, where sialyltransferases are normally located, in the isolated nerve terminals, a modulation of sialic acids would require the enzymatic machinery to be present in the active zone at the plasma membrane, implying a completely new mechanism for dynamic regulation of *N*-linked sialylation.

### Modulation of sialylation upon brief depolarization of synaptosomes

By using TMT reporter ion quantitation, the abundances of formerly sialylated *N-*linked glycopeptides in synaptosomes, stimulated with high KCl or a control buffer, were compared. Since protein translocation between the SDS-insoluble and SDS-soluble fraction was observed to a smaller extent, abundances were adjusted to the expression of the respective non-modified protein. However, this adjustment to the non-modified counterpart usually hinders the analysis, because membrane proteins of low abundance are often exclusively identified in the enriched sialylated fraction and not identified in the non-modified peptide fractions. To overcome these issues and increase the number of adjustable glycopeptides, all MS data from the non-modified peptide fractions were searched against two different databases. The first search included the whole taxonomy reference database, whereas the second database search was performed using a customized database created of all identified sialylated glycoproteins. As the reduction of the protein database could lead to too low estimates of the false discovery rate (FDR), the results of both searches were further evaluated in order to secure the quality of identification and quantitation (Figure S4). The second search identified 162 additional proteins that were not identified confidently in the first search (Figure S4A). This enabled the adjustment of 258 additional formerly sialylated *N-*linked glycopeptides in the SDS-insoluble fraction. These improvements were among others linked to the increased number of PSMs identified per protein when searching with the reduced glycoprotein database (Figure S4B). Comparing the correlations of all identified proteins using the Swissprot mouse/rat database search or the reduced glycoprotein database search showed that the results obtained with a reduced database gave higher correlations not only among replicates but also among conditions (Figure S4C). To validate the quantitation of newly identified glycoproteins, the quantitation of the same set of proteins either quantified by using the Swissprot mouse/rat or the reduced glycoprotein database were compared. Pearson correlations showed a higher correlation among replicates and among conditions when the search was performed with a reduced glycoprotein database. These results supported the use of a second MS database search of the non-modified dataset to increase the number of formerly sialylated *N-*linked glycopeptides being adjusted to the protein expression level. However, even with this strategy 35 % of all sialylated glycopeptides in the SDS-insoluble fraction could not be adjusted due to the lack of their non-modified counterpart.

Since a total of 9.4 % or 16.0 % of the formerly sialylated *N*-linked glycopeptides from the SDS-insoluble or SDS-soluble fraction could be adjusted and showed a significant change in sialylation upon depolarization, we expect to see the same percentage of significantly changing peptides for formerly sialylated *N*-linked glycopeptides that could not be adjusted to the protein level. Indeed, statistical testing of not adjustable formerly sialylated *N-*linked glycopeptides showed a similar ratio (15.8 - 16.9 %) compared to the adjusted ones. Therefore, both adjusted and not adjusted glycopeptides were used for further analyses.

Brief depolarization of synaptosomes resulted in the identification of a total of 228 significantly changing formerly sialylated *N-*linked glycopeptides in the SDS-insoluble fraction, whereof 76 glycopeptides could be adjusted to the level of their respective protein. In the SDS-soluble fraction, 271 formerly sialylated *N-*linked glycopeptides were significantly changing, whereof 156 glycopeptides could be adjusted to the protein level. Only about 8 % of the significantly changing formerly sialylated *N-*linked glycopeptides were common in both fractions. This clearly shows that sialylation is a highly dynamic PTM that can be modulated during synaptic transmission similarly to phosphorylation and therefore could have significant influence on the synaptic transmission processes in nerve terminals.

Dynamic sialylation was mapped to proteins functioning as neurotransmitter receptors (glutamate and GABA receptors), ion channels and transporters (e.g. voltage dependent calcium channels, sodium/potassium transporting ATPases, potassium voltage-gated channels, sodium/potassium/calcium exchangers), synaptic vesicle proteins (e.g. synaptophysin, synaptoporin) or cell adhesion molecules (e.g. contactins, neural cell adhesion molecules, neuroligin) (Figure 3). Desialylation was found to be more abundant than sialylation. The SDS-insoluble fraction showed 189 formerly sialylated *N-*linked glycopeptides significantly decreasing after five seconds depolarization and 39 significantly increasing. In the SDS-soluble fraction, the abundances of 167 formerly sialylated *N-*linked glycopeptides were significantly reduced, while 104 deglycosylated peptides increased in their levels. Since TiO_2_ is 95 % selective towards sialylated glycopeptides^41,51^, it can be assumed that a change in the abundance of a deglycopeptide is linked to change of the sialic acid content of the *N-*linked glycans attached to the specific *N-*linked sites. Hence, desialylation seems to be more pronounced than sialylation of *N-*linked glycoproteins upon five seconds depolarization.

**Figure 3:**
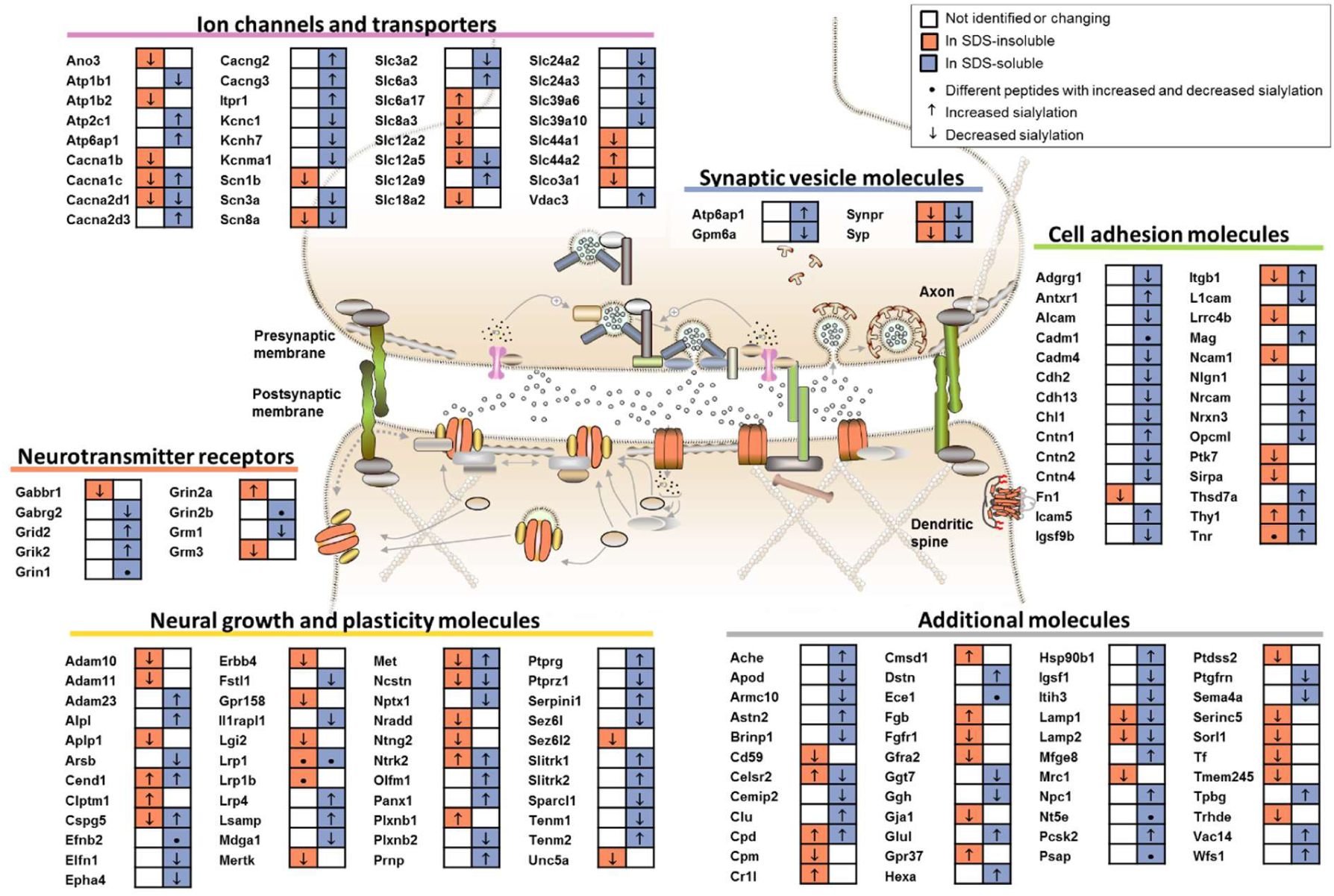
Examples of significantly changing formerly sialylated *N*-linked glycopeptides belonging to proteins essential for the synapse. Proteins showing a significant increase or decrease in sialylated glycosylation (LIMMA-testing q<0.05; only peptides adjusted to non-modified protein expression) upon synaptic transmission were classified into different groups regarding their function in the synapse. Significant changes of proteins from the SDS-insoluble and the SDS-soluble fraction are compared.

The biggest changes were observed for the Cell adhesion molecule 1 (Cadm1) increasing 4.2-fold in sialylated glycosylation at Asn-168 in the SDS-soluble fraction upon depolarization. Even though Cadm1 can carry polysialic acids, this site is not known to be affected with this modification^64^. In contrast, the sodium channel protein type 3 subunit alpha (Scn3a, Asn-1331) and the ionotropic glutamate receptor (Grin1, Asn-300) decreased in their sialylated glycosylation 2.6 and 2-fold, respectively. In the SDS-insoluble fraction, the highest changes were observed for the Sialomucin core protein 24 (Cd164, Asn-97) decreasing 2.8-fold and the Prenylcysteine oxidase (Pcyox1, Asn-196) increasing 1.4-fold in the abundance of formerly sialylated *N-*linked glycopeptides. Some proteins showed several peptides changing significantly in formerly sialylated *N-*linked glycosylation upon synaptic transmission (Figure S5). For instance, we identified eight sialylated *N-*linked glycosites of the tyrosine-protein phosphatase non-receptor type substrate 1 (Sirpa) in the SDS-insoluble fraction, whereof four sites were significantly altered in sialylation upon synaptic transmission (Figure S6). Similarly, the lysosome-associated membrane glycoprotein 1 had four sialylated *N-*linked glycosites in the SDS-soluble fraction, but only two of them were significantly desialylated during depolarization. This indicates, that changes in sialylation are rather site-specific and do not follow the general protein expression.

The observed changes in sialylation, propose a new highly dynamic function of sialylation of *N-*linked glycoproteins in nerve terminals. So far, sialylation has mainly been seen as an essential but rather static modification important for brain structure and function^25^, brain development^26^ and axon guidance^65^. Therefore, many studies have focused on the most frequent glycoconjugates in neurons, the gangliosides, or on polysialic acids attached to glycoproteins^21^. Here, major changes in sialylation of essential synaptic proteins were detected within a very short time frame by enriching for sialylated *N-*linked glycopeptides using TiO2 chromatography^41^. Sialylated glycosylation might modulate signalling as it is known for other PTMs. Since synaptosomes have not previously been shown to contain ER or a Golgi apparatus and only low amount of protein is synthesized in the nerve terminal^66^, the changes in sialylated *N-*linked glycosylation give evidence to the fact that sialylation, apart from the classical pathways, can take place at the plasma membrane. This implies that sialyltransferases, similar to sialidases, could be localized to the plasma membrane in the nerve terminal.

Recently, an increase in sialidase activity at hippocampal plasma membranes within a few seconds was detected upon neural excitation by using rat brain slices and a histochemical imaging probe for sialidase activity^67^. This increase in sialidase activity, also shown by the increase in extracellular free sialic acids after 30 min, was linked to a change in the subcellular localization of sialidases, where they suggested that mainly Neu4 is responsible for the desialylation^67^. The increased sialidase activity was further shown to influence intracellular calcium levels and down-regulate glutamate release, thereby suggesting that sialidase functions as a negative feedback regulator upon synaptic transmission^68^. In contrast to sialidase activity reported at the plasma membrane of neurons, extracellular sialyltransferases were only reported on blood cells and cancer cells to our knowledge^69^. In these cell types, extracellular sialyltransferases, being capable of performing extrinsic sialylation, suggested that extracellular glycans can be modulated with respect to sialic acids^70,71^. In addition to this, it has been shown for platelets, that they can release sugar nucleotides functioning as a substrate for sialyltransferases^72^. For nerve terminals, the mechanism of protein sialylation is relevant, considering the observed changes in formerly sialylated *N-*linked glycosylation of proteins highly important for synaptic transmission.

### Identified sialidases and sialyltransferases in synaptosomes

In order to identify potential enzymes responsible for sialylation and desialylation, the proteomics data were screened for sialidases and sialyltransferases. Out of 17 different peptides belonging to sialyltransferases or sialidases, five peptides were found in the non-modified fractions, whereas the remaining were identified in the formerly sialylated *N-*linked glycopeptide fraction (Table 1). Apart from the α2-8-sialyltransferase St8sia3, all proteins were identified with one or two peptides. However, most of them were observed in different independent experiments or in the SDS-insoluble as well as SDS-soluble fraction, indicating the low abundance of many of these enzymes in synaptosomes. The identified sialidases (Neu1, Neu2, Neu4) are known to modify glycoproteins and previous reports have shown that at least Neu1 and Neu2 can be active at the plasma membrane under certain circumstances^73,74^. How these enzymes are associated with the membrane remains controversial. For Neu1, some studies suggested a C-terminal transmembrane domain, whereas other studies excluded a transmembrane region proposing a lipid anchor^75,76^. Subcellular location and activity of sialidases might differ depending on the physiological conditions and the cell types, as demonstrated by the polysialic acid processing by Neu1^77^ and the involvement of Neu4 in hippocampal memory processing^78^.

**Table 1:**
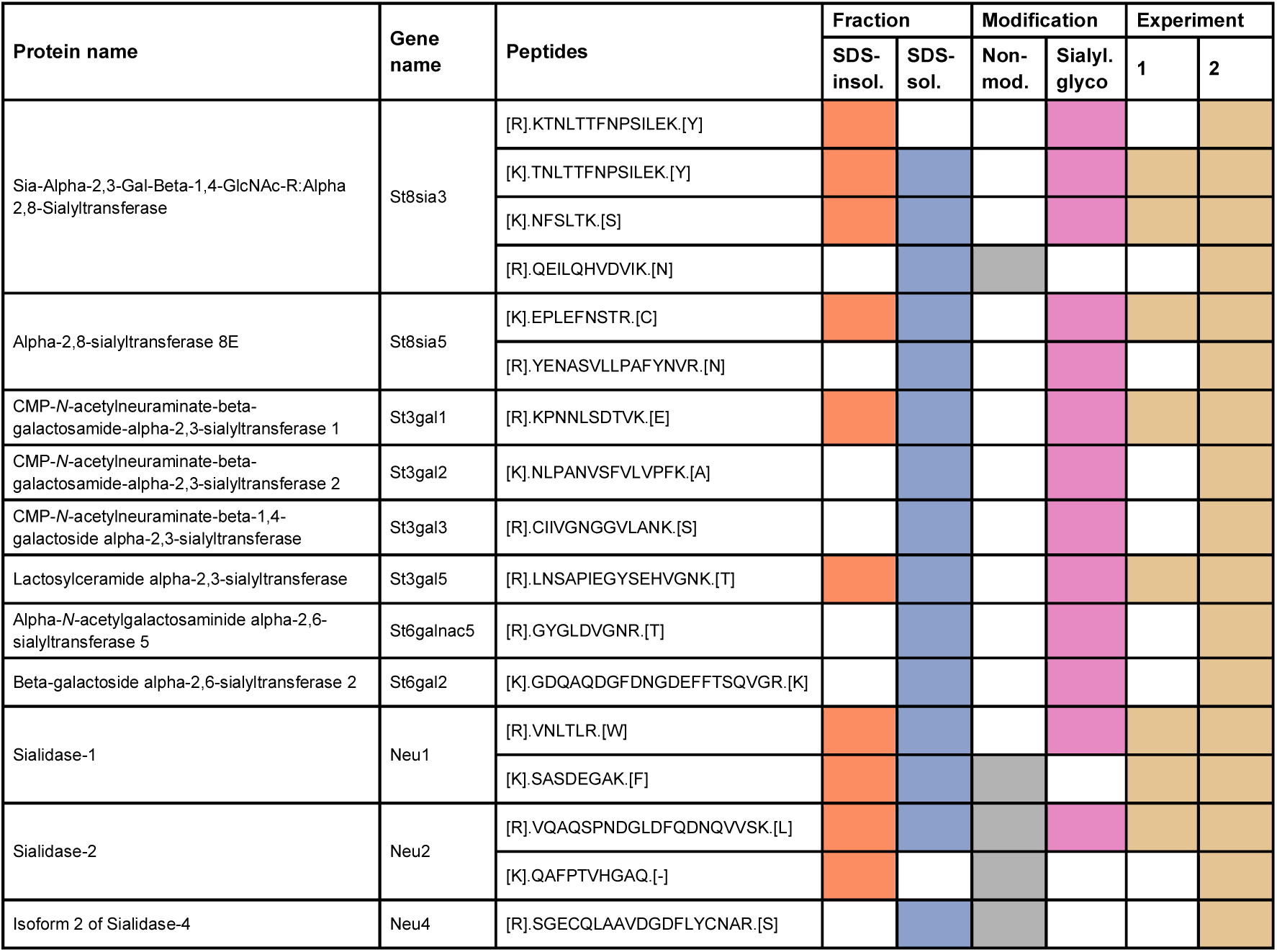
Sialidases and sialyltransferases identified in the SDS-insoluble and SDS-soluble fraction of synaptosomes. The peptides of sialidases and sialyltransferases identified in two proteomics experiments are listed. The colours encode for an identification in a certain fraction, modification group or experiment.

Regarding sialyltransferases, enzymes responsible for alpha-2,3, alpha-2,6 and alpha-2,8 sialic acid linkages were identified. Especially the sialyltransferases St8sia3 and St8sia5 have high potential in the modulation of sialylation in nerve terminals, since they can connect several sialic acids in an alpha-2,8 linkage producing among others disialic acid epitopes. Even though singly sialylated structures are common, a frequent occurrence of alpha-2,8-linked disialic and oligosialic acids in mammalian brain glycoproteins has previously been described^79^. This diSia epitope was shown to have a function in neurite formation and it was suggested that the sialyltransferase St8sia3 might be responsible for the synthesis of this epitope^80^. In order to understand the observed changes in sialylation upon synaptic transmission, the questions which sialyltransferases are active at the plasma membrane of nerve terminals, and which sialic acid linkages are preferably changing in sialylated *N-*linked glycosylation upon depolarization need to be addressed.

Since the MS data only provided the existence of sialidases and sialyltransferases, further investigations regarding their activity in synaptosomes are needed. In addition to sialyltransferases, β1-3 and β1-4 galactosyltransferases as well as UDP-GalNAc:beta-1,3-*N-* acetylgalactosaminyl-transferase and Beta-1,4 *N-*acetylgalactosaminyl-transferases were identified in the dataset. It remains to be determined, if these enzymes were located at the synaptosomal membrane, if they are present in other vesicle-like organelles such as endosomes, or if they represent contaminants. Alternatively, traces of Golgi could be present at the nerve terminal.

### Potential effects of modulation of sialylated N-linked glycosylation in the postsynaptic density and synaptic vesicles (SVs)

Interestingly, many proteins that can be in the postsynaptic membrane showed changes in sialylated glycosylation (Figure 4A). Of these, voltage gated ion channels, ionotropic glutamate receptors, transmembrane receptors and cell adhesion molecules had common downstream interactors such as calcium/calmodulin dependent protein kinases, cAMP-dependent protein kinases, Src kinases, Ras proteins and disks large homolog proteins including PSD-95. This direct link to protein kinases indicates an interconnection of the two different PTMs. Since synaptosomes are not metabolically active in their postsynaptic density due to the lack of mitochondria and ATP, changes in phosphorylation on the postsynapse side cannot be determined using this model system. However, it is highly interesting to investigate, if changes in sialylated *N-*linked glycosylation are altering protein phosphorylation, as we previously have shown for another excitable cell type, pancreatic β-cells during glucose assisted depolarization and insulin release^81^. The downstream interactors suggested a link between protein sialylation and long-term potentiation (LTP), endocytosis, exocytosis and hippo pathway signalling just to name a few biological processes. Focusing on LTP, not only downstream interactors were involved in this process. Significantly altered sialylated glycoproteins such as Grin2a, Grin2b, Nlgn1, Ntrk2, Slc8a3, Slc24a2 and Tnr are directly involved in LTP^82–87^. This suggests, that sialylated glycosylation might also play a role in synaptic plasticity and memory formation given the fact that the five seconds stimulation exceeded by far the duration of a single transmission.

**Figure 4:**
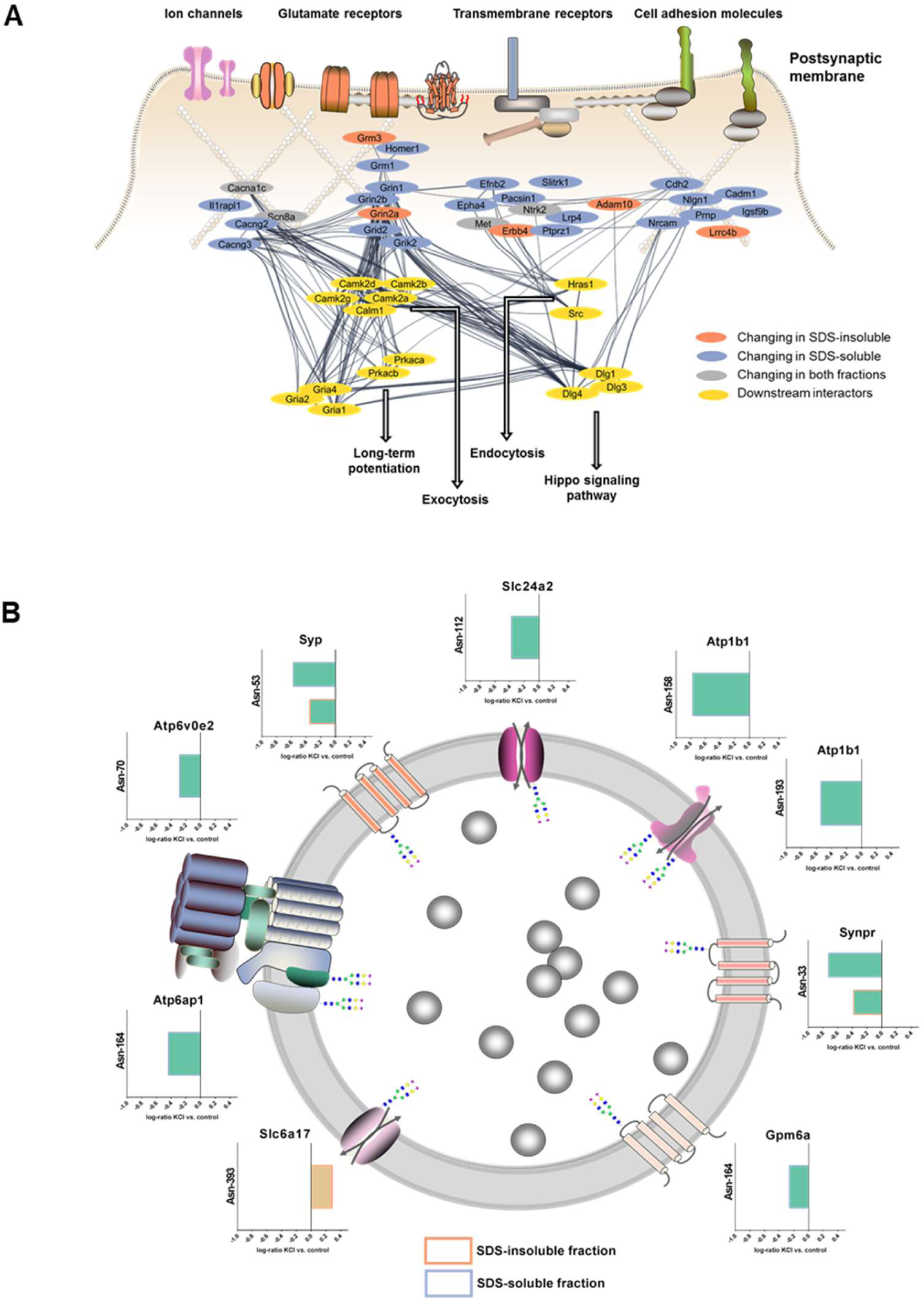
Effects of changes in sialylated *N-*linked glycosylation on signalling in the postsynaptic density and synaptic vesicle function. A) Postsynaptic proteins showing significant changes in sialylated *N-*linked glycosylation upon depolarization were used to create a protein-protein interaction network. Downstream interactors and signalling pathways were identified by allowing 15 additional interacting proteins that were not significantly changing or detected in the dataset. B) Log-ratios (KCl vs. control) of the significantly changing sialylated synaptic vesicle proteins, Glycoprotein M6a (Gpm6a), Sodium-dependent neutral amino acid transporter (Slc6a17), Sodium/potassium/calcium exchanger 2 (Slc24a2), Sodium/potassium-transporting ATPase subunit beta-1 (Atp1b1), Synaptophysin (Syp), Synaptoporin (Sypr), V-type proton ATPase subunit e2 (Atp6v0e2) and V-type proton ATPase subunit S1 (Atp6ap1) are illustrated.

In addition to proteins in the postsynaptic membrane, we identified significant changes in sialylated glycosylation of synaptic vesicle proteins identified in the proteomics map of the synaptic vesicle presented by Takamori *et al*.^14^ (Figure 4B). In particular, the glycoprotein M6a (Gpm6a), the sodium/potassium/calcium exchanger 2 (Slc24a2), the sodium/potassium-transporting ATPase subunit beta-1 (Atp1b1), synaptophysin (Syp), synaptoporin (Synpr), the V-type proton ATPase subunit e2 (Atp6v0e2) and V-type proton ATPase subunit S1 (Atp6ap1) were desialylated after brief depolarization with KCl. Only the sodium-dependent neutral amino acid transporter (Slc6a17) showed an increased abundance of a formerly sialylated *N-* linked glycopeptide upon synaptic transmission. As shown for synaptophysin (Syp) and subunits of the V-type ATPase (Atp6v0b2) in particular, glycosylation is often associated with protein stability and correct localization^88,89^. For some transporters similar to those determined in this study, *N-*glycan removal led to a reduced activity^90^.

Even though the expression of sialidase in synaptic junctions was revealed, a clear evidence for sialidase localization in synaptic vesicles has not been given, yet^30,91^. Proteomics analysis of synaptic vesicles did neither observe sialidases nor sialyltransferases^14,92^, nontheless these studies did not enrich for low abundant proteins. Nevertheless, we observed desialylation of a subunit of the vacuolar ATPase, which acidifies synaptic vesicles to enable the filling of SVs with neurotransmitters and is only present in one or two copies per vesicles^14^. The E2 subunit of the V0 sector which was desialylated at Asn-70 is expressed only in restricted tissues including the brain^93^. Nothing is known about glycosylation of this subunit in particular, albeit glycosylation of lysosomal v-ATPase was previously shown to influence proton pump activity and signalling in fibrosis^94^. Furthermore, the accessory subunit Ac45 (Atp6ap1) was observed to decrease in sialylated glycosylation at Asn-164 after depolarization and this protein is known to influence v-ATPase function^95^. Apart from ion pumps and transporters, synaptophysin and synaptoporin, two homologous proteins, were both desialylated at the same site (Syp Asn-53, Synpr Asn-33) which was observed in the SDS-insoluble as well as SDS-soluble fraction. The role of synaptophysin and synaptoporin in synaptic vesicles, where they represent about 10 % of the total protein content^14^, is still not completely clear, but it has been revealed that synaptophysin contributes to efficient endocytosis^96^. Synaptophysin’s *N-*linked glycosylation site Asn-53 was shown to be essential for localization in synaptic vesicle membranes and recycling in response to neural activity^88^. Generally, sialylation of synaptic vesicle proteins might contribute to an intravesicular proteoglycan matrix^97^. Reigada *et al*. proposed that negatively charged residues of keratan sulphate and sialylated glycoproteins interact with neurotransmitters such as acetylcholine and thereby create a smart gel within synaptic vesicles that controls neurotransmitter release^97^. Desialylation of synaptic vesicle proteins upon synaptic transmission might thereby increase neurotransmitter accessibility.

Taken together, essential synaptic vesicle molecules involved in neurotransmitter filling, ion exchange and endocytosis changed significantly in *N-*linked sialylation after depolarization indicating a role of sialic acids in mediating SV cycling, neurotransmitter release and potentially filling up the SVs with neurotransmitters.

### Potential molecular role of sialylation of the NMDA receptor molecule

To address the question, how changes in sialylated *N-*linked glycosylation could affect the signalling upon synaptic transmission, the location of significantly changing formerly sialylated *N-*linked glycosites on the cryo-EM structure of the GluN1-GluN2B NMDA receptor including the exon 5 was examined in detail (Figure 5)^98^. Five formerly sialylated *N-*linked glycosites that were changing during depolarization were mapped to the heterotetrameric NMDA receptor structure (GluN1/Grin1: Asn-321↓, Asn-389↓, Asn-461↑; GluN2B/Grin2b: Asn-348↑, Asn-444↓). The GluN1 subunit had one modified glycosylation site (Asn-461) in the ligand binding domain facing the outside of the receptor which showed an increase in sialylation after depolarization. Two additional *N-*linked glycosites decreasing in sialylation upon synaptic transmission were located in the amino-terminal domain (ATD) (Asn-321, Asn-389). Interestingly, the cryo-EM protein structure indicated that one *N-*linked glycosite (Asn-321) is positioned at the apex of the NMDA receptor in the upper lobe of the ATD. Binding of ligands such as zinc and changes of amino acid interactions in the upper lobe were shown to affect the conformation of NMDA receptors and the opening and closing of the channel^99^. However, to our knowledge the specific function of this region of the GluN1 subunit including Asn-321 is unknown. Due to the pairwise identical subunits, two sialylated *N-*linked glycans of the Asn-321 site are in proximity at the top of the receptor complex. Even though the protein was expressed in insect cells and the shown glycans might not represent the ones present in rat nerve terminals, changes in sialylation are likely to affect the interaction of the two subunits and modulate protein structure and function at the top of the receptor complex. In this case, desialylation occurred during five seconds of depolarization, meaning that negative charges were removed and that might result in less repulsion of the two glycan structures and affect the gating of the ion channel^100^. As mentioned above, glycans are essential for the structure and function of ion channels. Glycans were observed to decrease the function of pentameric ligand gated ion channels and they affect the gating properties of potassium channels^34,101^. For the GluN1, GluN2B NMDA receptor, it was predicted that glycans in the ligand binding domain stabilize the closed conformation^102^. However, the effect of changes in sialylation at specific *N-*linked glycosites and on distinct glycan structures on the function of ion channels remains to be investigated. In addition, since sialic acid is negatively charged at physiological pH it is fair to speculate that increased sialylation on Asn-321 of the GluN1 subunit could result in ion channel opening functions allowing transportation of ions in and out of the receptor ion channel.

**Figure 5:**
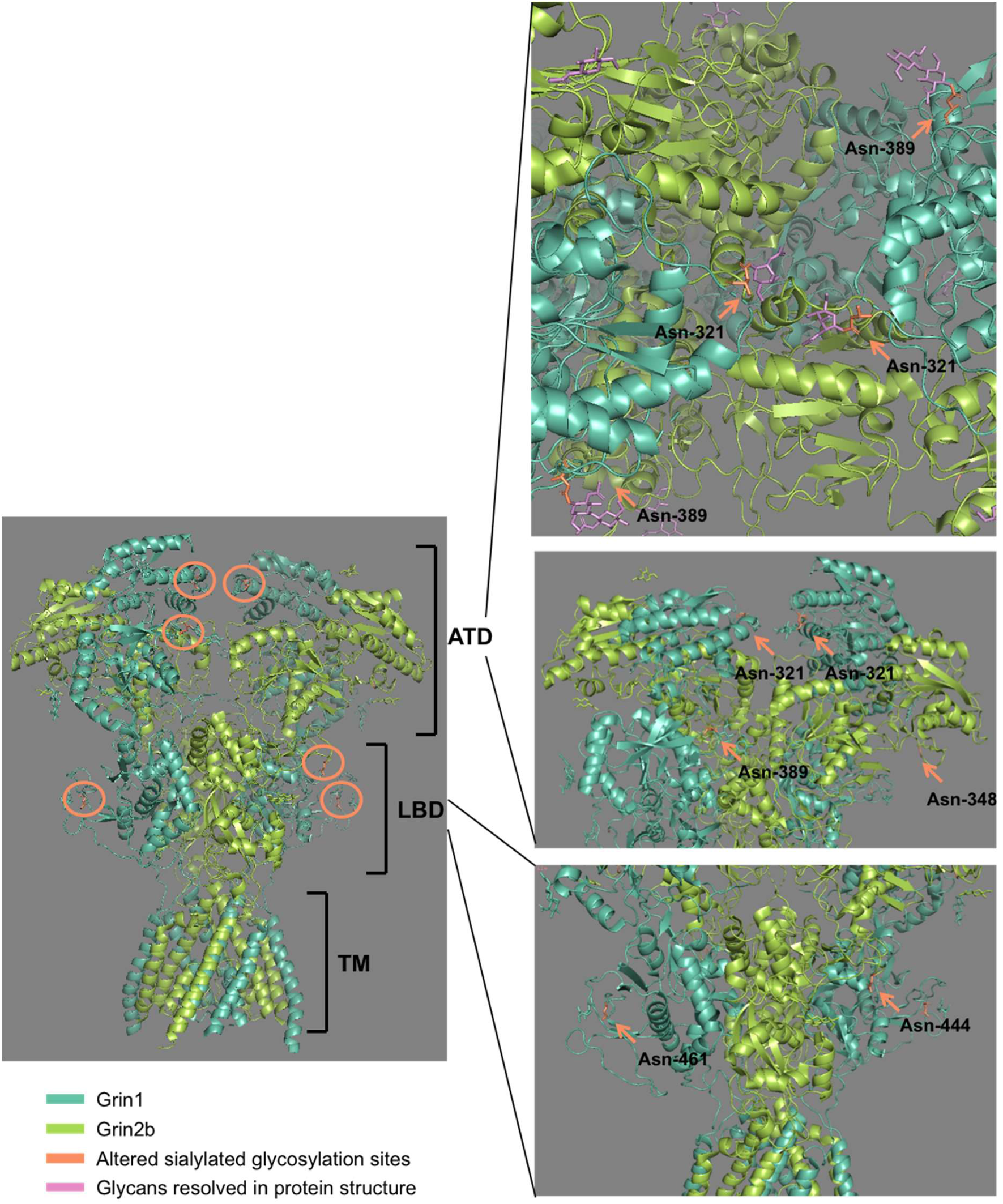
Molecular structure of the heterotetrameric GluN1-GluN2B NMDA receptor and altered sialylated *N-*linked glycosites. The cryoEM structure of the rat GluN1-GluN2B NMDA receptor with exon 5 expressed in insect cells was downloaded from the RCBS protein data bank and all formerly sialylated *N-*linked glycosites that were significantly changing upon depolarization were highlighted in orange. The receptor consists of an amino-terminal domain (ATD), a ligand binding domain (LBD) and a transmembrane domain (TM).

Furthermore, it has recently been suggested that X-ray structures of the Cys-loop receptors, such as the GABA_A_ receptor, may reflect a desensitized state103. Many high-resolution glycoprotein structures have the drawback that glycans needed to be removed in order to allow crystallization and subsequent analysis of the protein structure. In addition to this, mammalian proteins are often expressed in non-mammalian expression systems resulting in less complex glycan structures.

So far, the function of modulation of sialylated glycosylation upon synaptic transmission can only be estimated due to the lack of molecular tools for characterizing or manipulating site-specific sialylation. Site-directed mutagenesis is not sufficient, since it removes the whole *N-* linked glycan and the various glycan compositions, which could result in misfolding of the protein and the risk of degradation in the proteasome or significant reduction in activity. On the contrary, enzymatic removal of sialic acids from *N-*linked glycans targets only sialic acids, but it is not site-specific. Novel techniques allowing site and glycan-specific modulation remain to be developed in order to allow functional validation of sialylation and the study of this interesting PTM in biological systems.

## Conclusion

By isolating nerve terminals from rat brains and separating the proteins into an SDS-insoluble and an SDS-soluble fraction, a global roadmap of the synaptic proteome and *N-*linked sialiome was achieved using state-of-the-art enrichment of sialylated glycopeptides using TiO_2_ chromatography in combination with enzymatic deglycosylation and LC-MS/MS. In addition, this study showed for the first time the dynamics of sialylated *N-*linked glycosylation on synaptic proteins upon very brief depolarization of nerve terminals. The addition or removal of negative charges by sialylation or desialylation could facilitate the signalling process of synaptic transmission in a global manner and influence down-stream phosphorylation dependent processes similarly to what we previously have shown for glucose stimulated insulin secretion in pancreatic β-cells^81^. We identified potential sialidases and sialyltransferases that might be responsible for these changes, though further investigation is needed to identify the exact mechanism of sialylation/desialylation at the synaptic membrane. In contrast to phosphorylation being attached directly to the polypeptide backbone, sialic acids are linked to complex glycan structures exhibiting a huge microheterogeneity. Furthermore, contrasting other PTMs such as phosphorylation, site-specific modulation of sialic acids on specific glycans and specific *N-*linked glycosites is not possible with present molecular tools. Understanding the preference of specific glycosites or glycan structures for dynamic changes in sialylation and assigning the specific enzyme responsible for the addition or removal of the sialic acids on specific sites are needed in the future to expose molecular functions to each dynamic change described in this study. The present study demonstrated for the first time a new regulatory level of synaptic transmission and provides a starting point for protein specific investigation, unravelling the role of site-specific sialylation dynamics in nerve terminals.

## Acknowledgements

This study was supported by the Danish Natural Science Research Council (grant number: 6108-00621B). The Villum Center for Bioanalytical Sciences at SDU is acknowledged for access to state-of-the art mass spectrometric instrumentation.

## Data availability

LC-MS/MS raw data and search results are deposited in the PRoteomics IDEntifications (PRIDE) database (https://www.ebi.ac.uk/pride/archive/) with the dataset identifier PXD016230 ^104^.

