## Supplemental Figures for "Synaptic transmission induces site-specific changes in sialylation on *N*-linked glycoproteins in rat nerve terminals"

### Supplemental data

**Figure S1:** Characterization of non-modified proteins identified in SDS-insoluble and SDS-soluble fractions

**Figure S2:** Evaluation of the depolarization of synaptosomes

**Figure S3:** Characterization of all identified sialylated *N*-linked glycoproteins by KEGG pathway and gene ontology (biological process) enrichment analysis.

**Figure S4:** MS data searching using a reduced glycoprotein database for the enhanced identification of glycoproteins in the non-modified dataset.

**Figure S5:** Protein-protein interaction networks and KEGG pathway enrichment of proteins with significantly changing sialylated *N*-linked glycosylation upon depolarization.

**Figure S6:** Site-specific changes of sialylated *N*-linked glycosylation on Sirpa and Lamp1.

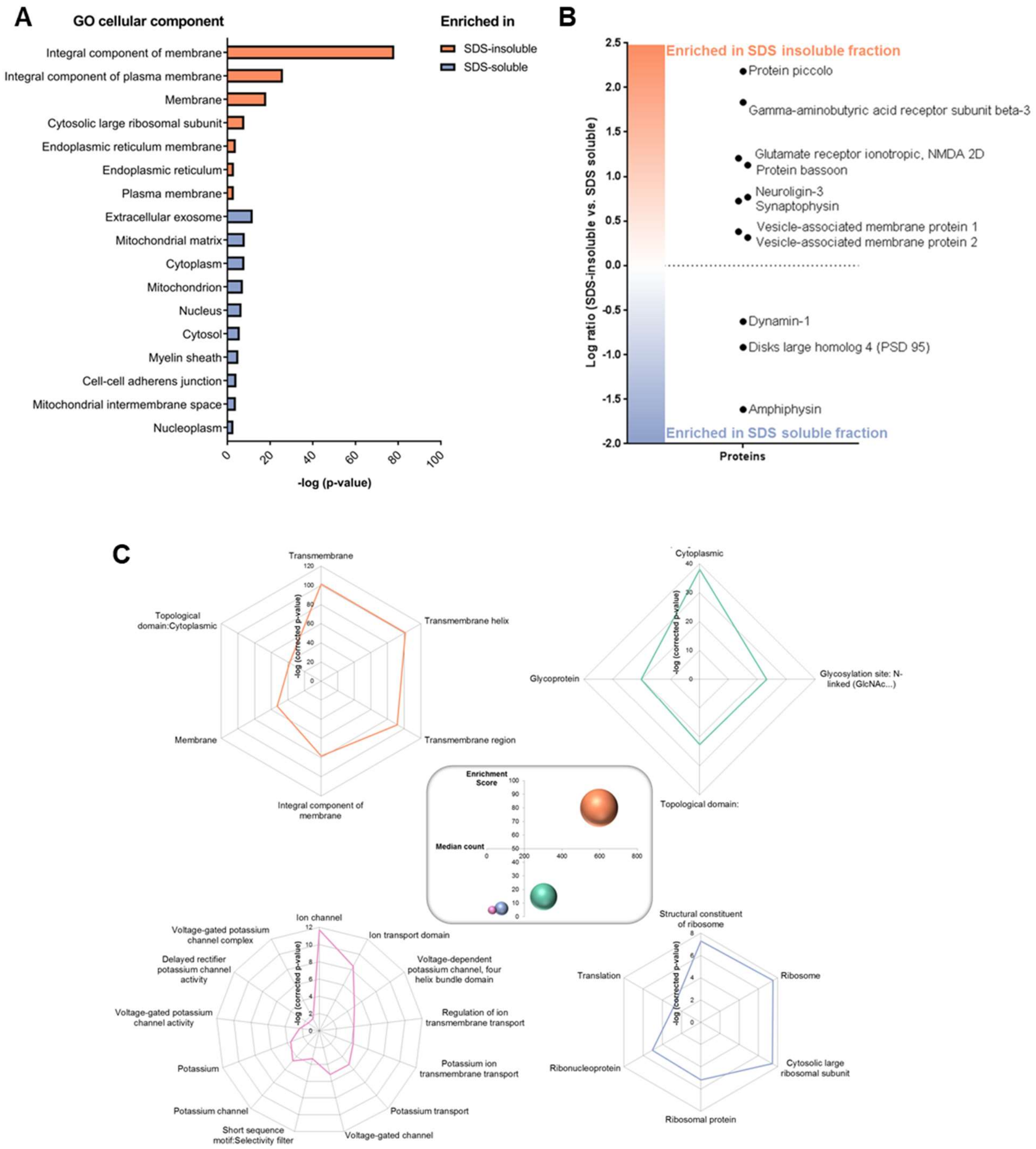


**Figure S1: Characterization of non-modified proteins identified in SDS-insoluble and SDS-soluble fractions.** (A) Gene ontology (cellular component) enrichment analysis was performed with proteins significantly enriched in the SDS-insoluble or SDS-soluble fraction (LIMMA testing q<0.05) using all identified proteins as a background. (B) Log-ratios (SDS-insoluble vs. SDS-soluble) of non-modified proteins that represent active zone, postsynaptic density or endocytosis markers were plotted. (C) For proteins that were significantly enriched in the SDS-insoluble fraction, functional annotation clusters were analysed. The enrichment score and the median count of proteins for all four clusters are given in the panel in the middle, whereas each cluster is further described by their individually enriched terms plotted with the –log p-value after Benjamini-Hochberg correction.

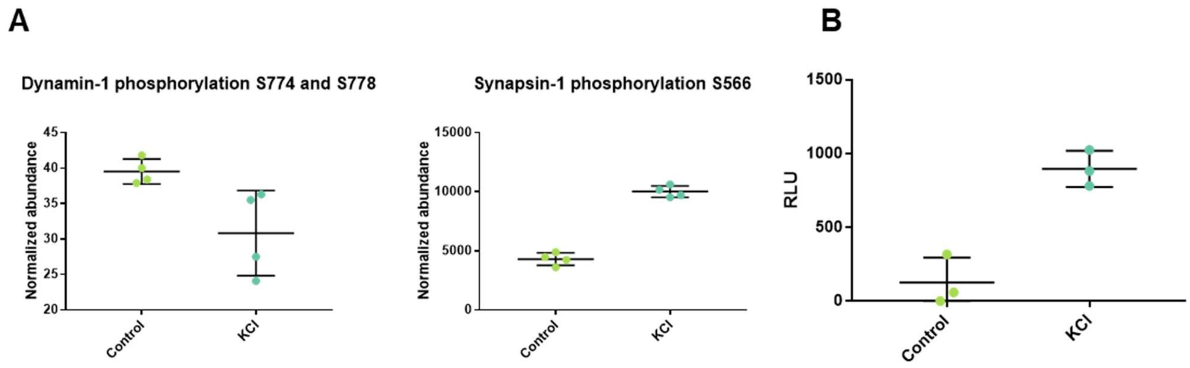


**Figure S2: Evaluation of the depolarization of synaptosomes.** (A) Phosphorylation and dephosphorylation of the marker proteins Dynamin-1 and Synapsin-1 were determined by quantitative proteomics. (B) The glutamate release from synaptosomes stimulated with a control HBK or a 76.2 mM KCl HBK buffer was measured by using the Glutamate Glo Assay determining the relative luminescence units (RLU).

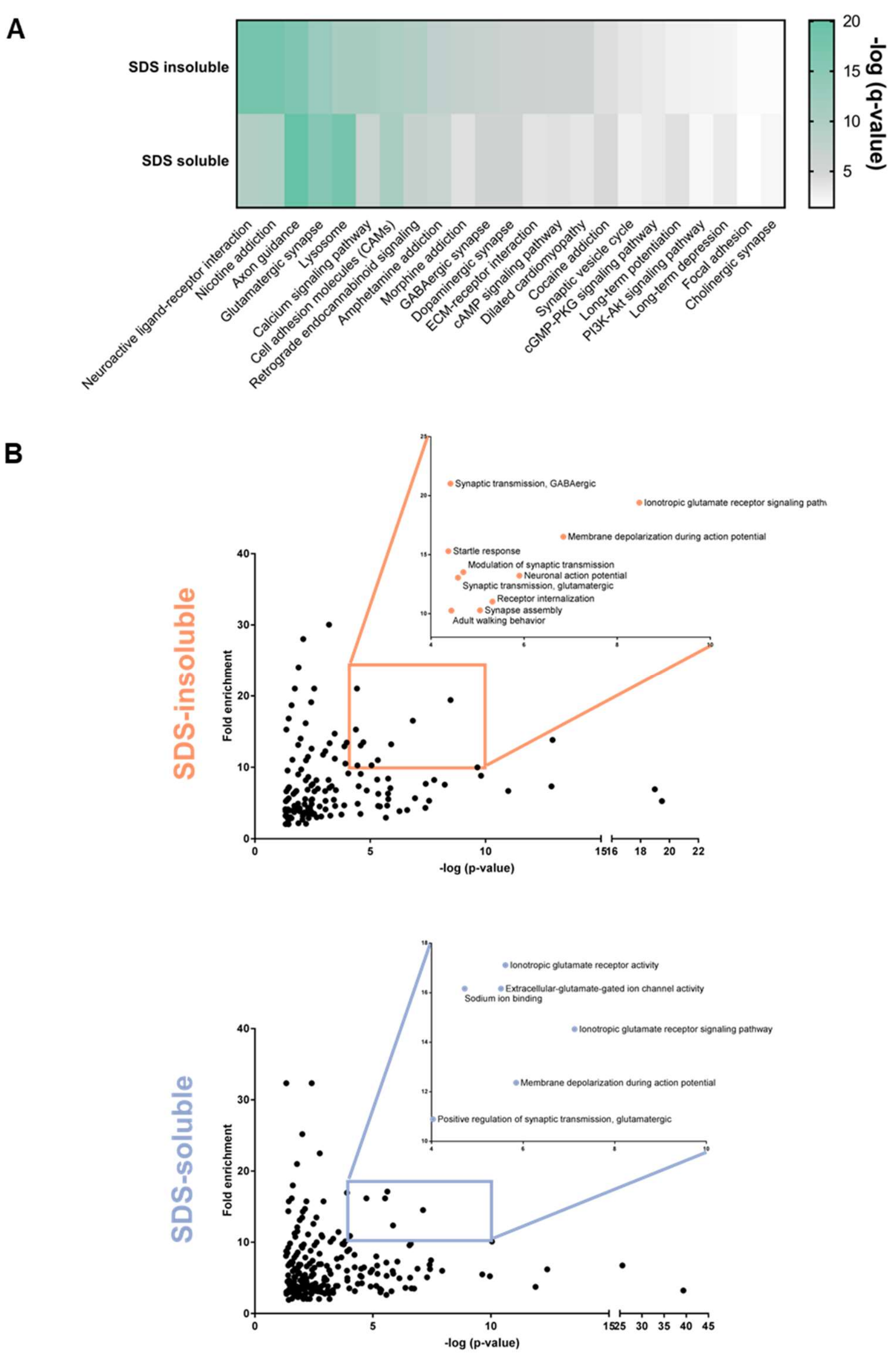


**Figure S3: Characterization of all identified sialylated *N*-linked glycoproteins by KEGG pathway and gene ontology (biological process) enrichment analysis.** (A) The enriched KEGG pathways of all identified sialylated glycoproteins of the SDS-insoluble and SDS-soluble fraction were compared (Benjamini-Hochberg corrected p-value < 0.05). (B) The most relevant KEGG pathways/ gene ontologies (Benjamini-Hochberg corrected p-value < 0.05) were extracted by plotting the fold enrichment against the –log p-value. Terms with a high fold enrichment and a high –log p-value were highlighted, and their terms were shown in detail.

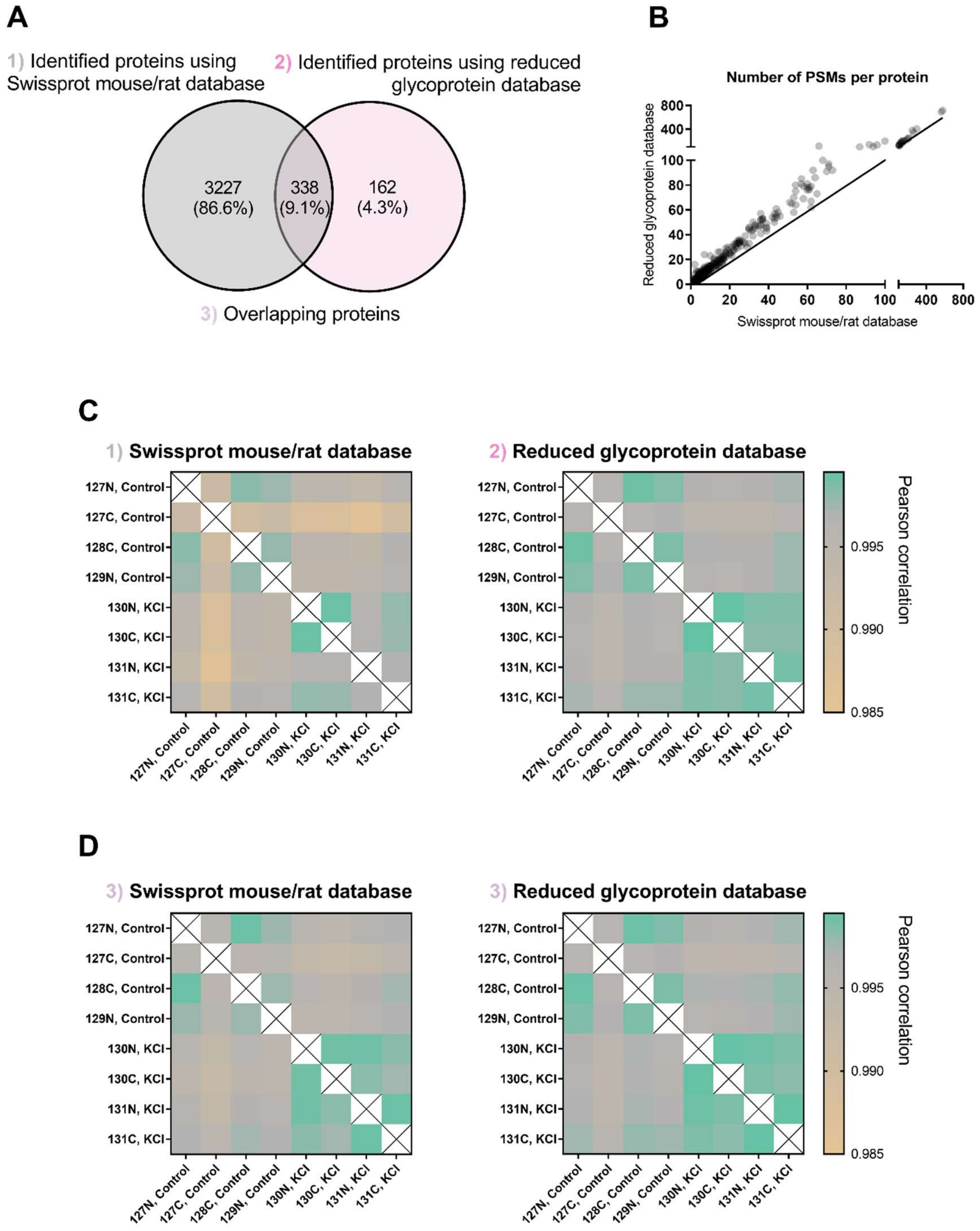


**Figure S4: MS data searching using a reduced glycoprotein database for the enhanced identification of glycoproteins in the non-modified dataset.** (A) Non-modified proteins of the SDS-insoluble fraction identified by using the entire Swissprot mouse/rat database or a reduced glycoprotein database for the MS data searching were compared. (B) The number of PSMs per protein identified using a reduced glycoprotein database was plotted against the number of PSMs per protein identified using the entire Swissprot mouse/rat database. The line represents an equal number of PSMs for one protein determined in both searches. (C) Pearson correlations of all identified proteins using the different databases were calculated. (D) Pearson correlations of proteins identified in both MS data searches but quantified differently were compared.

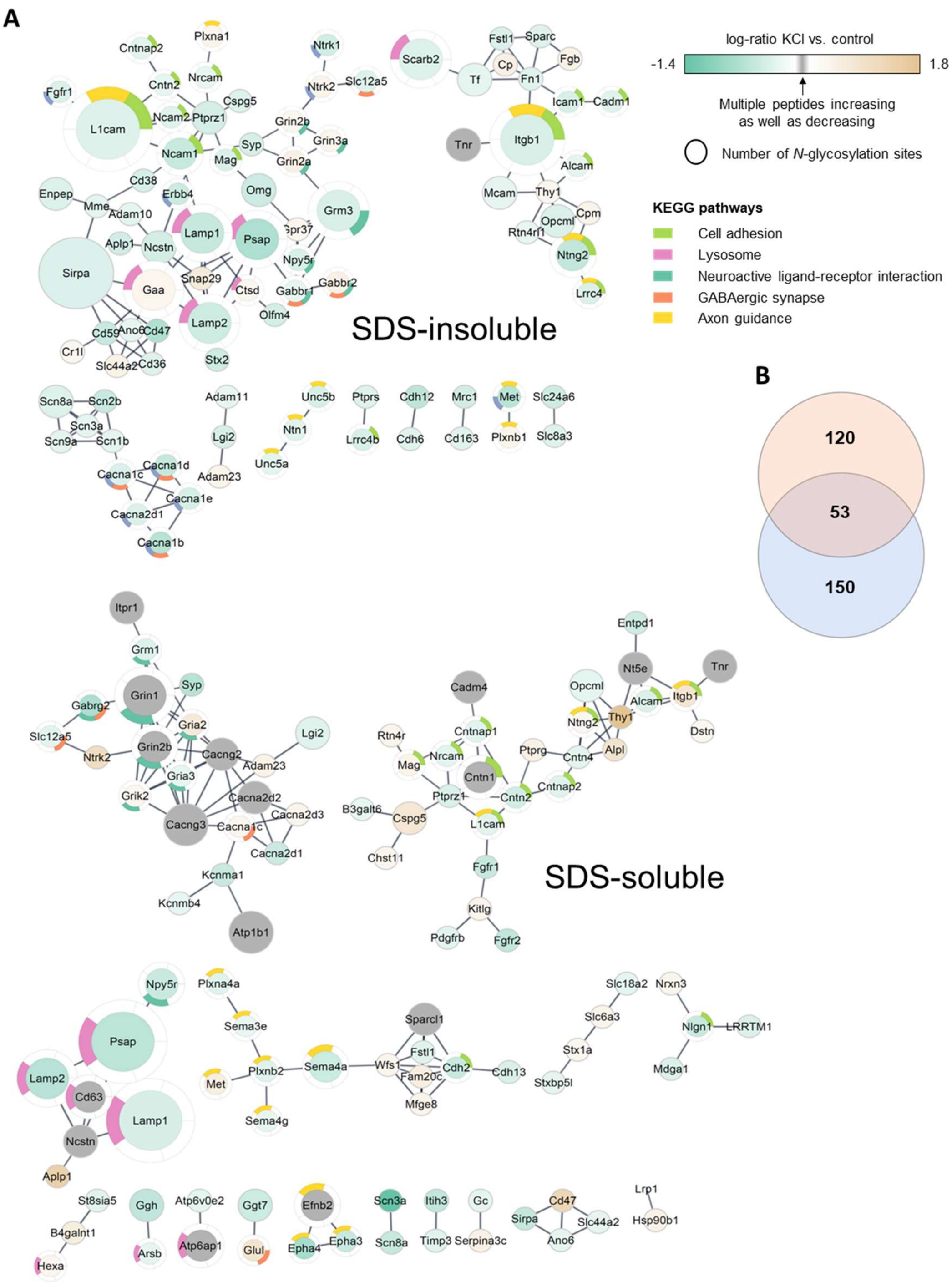


**Figure S5: Protein-protein interaction networks and KEGG pathway enrichment of proteins with significantly changing sialylated *N*-linked glycosylation upon depolarization.** (A) Significantly changing formerly sialylated *N*-linked glycopeptides (not adjusted and adjusted) from the SDS-insoluble and SDS-soluble fraction were rolled up to proteins and their protein-protein interactions as well as KEGG pathway enrichment were investigated by using STRING. Only connected nodes are shown. (B) Altered proteins from both fractions (SDS-insoluble = orange and SDS-soluble = blue) were compared in a Venn diagram.

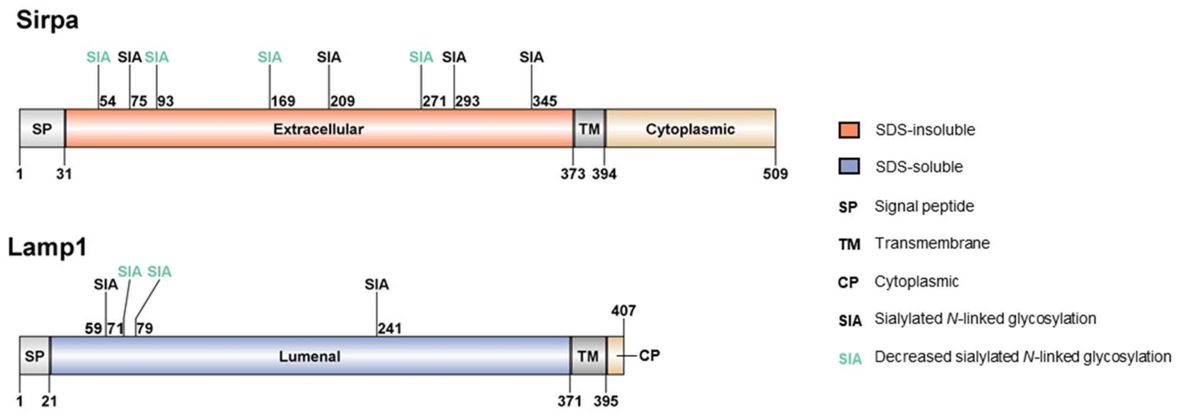


**Figure S6: Site-specific changes of sialylated *N*-linked glycosylation on Sirpa and Lamp1.** For the tyrosine-protein phosphatase non-receptor type substrate 1 (Sirpa, SDS-insoluble fraction) and the lysosome-associated membrane glycoprotein 1 (Lamp1, SDS-soluble fraction), all identified sialylated *N*-linked glycosites were mapped, whereas significantly desialylated sites were highlighted in green.
